# The impact of data reduction and lossy image formats on electron microscope images

**DOI:** 10.1101/451427

**Authors:** Edward T. Eng, Mykhailo Kopylov, Carl J. Negro, Sarkis Dallaykan, William J. Rice, Kelsey D. Jordan, Kotaro Kelley, Bridget Carragher, Clinton S. Potter

**Affiliations:** Simons Electron Microscopy Center, New York Structural Biology Center, 89 Convent Ave, New York, NY, 10027, USA.; Department of Biochemistry and Molecular Biophysics, Columbia University, New York, NY 10032, USA.

**Keywords:** electron microscopy, file format, image data, data archiving, structural biology

## Abstract

Recent advances in instrumentation and software for cryoEM have increased the applicability and utility of this method. Coupled with the adoption of automated pipelines, significant infrastructure support is required to sustain high throughput workflows. In particular, data generation rates may outpace the ability to deploy data storage and archival solutions. We have investigated what effects data compression and conversion to different file formats have on the ability to obtain high resolution cryoEM reconstructions. Standard lossless data compression strategies have a high impact on reducing the size of direct detector electron counting movie stacks, but provide more modest gains for aligned summed images. We show that EM images can be compressed using standard lossy methods to reduce file storage to 5–10% of the size of the original aligned sum or movie stack file and yet still retain enough information such that modern image processing pipelines will provide sub-2Å reconstructions from the compressed data.

## Introduction

The application of robust automated data acquisition systems and processing pipelines in the field of cryogenic electron microscopy (cryoEM) has increased throughput and the ability to apply this technique to a greater variety of structural biological applications (Baldwin et al., 2018). Along with advances in detector technology, data generation rates are also growing due to faster frame rates and larger sensor sizes. Meanwhile, the field is still in the process of developing standards as to what forms of the experimental data should be saved and available to the community, adding uncertainty to the scope of data that needs to be archived (Henderson et al., 2012, Patwardhan et al., 2016, Kleywegt et al., 2018).

A data retention and management policy for three dimensional EM has been recognized as an issue that is crucial for the field, scientific computing and future research (Patwardhan et al., 2012). Applying the next generation of computational software to existing data may lead to further gains in resolution and understanding of the underlying biology. In particular, direct detector frame stacks have been required for refinement improvements because of the developments of improved motion and dose correction programs (Ripstein and Rubenstein 2016). For example, several researchers have processed publicly available archived datasets to improve 3D reconstructed maps by ∼1Å (Tegunov et al., 2018 and Heymann et al., 2018).

A concern is that researchers may not have the resources to backup and archive all the data generated over the course of an experiment, and instead may focus on the minimal set that was used for the published reconstructed map. Use of publicly available resources, such as Electron Microscopy Public Image Archive (EMPIAR), Electron Microscopy Data Bank (EMDB) and Protein Data Bank (PDB), does not obviate the researcher’s responsibility for archival backup. Data reduction and compression are important strategies used throughout computing, but for cryoEM structural biology, the primary goal is a reduction in the file size to conserve data storage resources. A robust data reduction strategy can extend the lifetime of existing computational infrastructure and increase capacity for data retention. It is notable that more packages are now supporting compressed TIFF formats throughput the processing pipeline (Grant et al., 2018 and Zivanov et al., 2018).

An important factor in deciding best practices is to understand the tradeoffs of the current options in the field. Here, we describe the potential disk usage savings gained by storing data using various common file formats and compression standards. We also analyze the effect on cryoEM reconstructions if the full data processing pipeline is applied to compressed data. While the preference may be to always retain experimental data with complete fidelity, we have determined that a significant amount of data loss can be tolerated and still retain the ability to reproduce experimental results.

## Material and Methods

The apoferritin datasets used for this study were downloaded from EMPIAR (EMPIAR-10200 and EMPIAR-10146). The bovine glutamate dehydrogenase (GDH) and *Thermoplasma acidophilum* 20S (T20S) datasets were collected in house.

### Samples

GDH was purchased from Sigma (G2626) and purified by anion exchange chromatography, followed by size-exclusion chromatography in 150 mM NaOH, 20mM HEPES, pH 7.4. T20S proteasome was a gift from Yifan Cheng, Zanlin Yu, and Kiyoshi Egami. The received stock was separated into small aliquots and stored at −80 °C in 20 mM Tris, pH 8.0, 50 mM NaCl, 0.1 mM EDTA.

### Grid Preparation

3 µl of freshly thawed protein (4 mg/mL for GDH and 0.29 mg/mL for T20S) was applied to plasma-cleaned C-flat 1.2/1.3 400 mesh Cu holey carbon grids (Protochips, Raleigh, North Carolina), blotted for 2.5 s after a 30 s wait time, and then plunge frozen in liquid ethane, cooled by liquid nitrogen, using the Cryoplunge 3 (Gatan) at 75% relative humidity.

### Microscopy

Thermo-Fisher Titan Krios operated at 300 kV and Gatan K2 Summit camera in counting mode were used with a 70 µm C2 aperture, 100 µm objective aperture and calibrated pixel size of 0.6616Å.

### Imaging

Movies were collected in counting mode using Leginon (Suloway et al., 2005). For the T20S sample, a dose rate of 6 e^−^/Å^2^/s was used for an accumulated dose of 62.59 e^−^/Å^2^ with a total exposure time of 10 seconds and 0.2 second frame rate. For the GDH sample, a dose rate of 10 e^−^/Å^2^/s was used for an accumulated dose of 61.25 e^−^/Å^2^ with a total exposure time of 6 seconds and 0.1 second frame rate. Nominal defocus values ranged from 1.0–3.0 µm.

### Image Processing

Movies recorded on the K2 were aligned using MotionCor2 with dose weighting (Zheng et al., 2017) and CTF estimation was performed with CTFFIND4 (Rohou and Grigorieff, 2015). For the first 50 images particles were picked automatically using the Appion DoG Picker (Voss et al., 2009), extracted, and subjected to 2D classification in RELION to create templates for another round of particle picking using FindEM (Roseman, 2004). The picked particles were extracted and subjected to 2D classification and the best classes were selected for *ab-initio* reconstruction to create an initial model in cryoSPARC (Punjani et al., 2017). This model was used for 3D classification in RELION or heterogeneous refinement in cryoSPARC. For the final reconstruction, particles were selected and subjected to 3D refinement in cryoSPARC, RELION or cisTEM. For the T20S reconstruction 1173 micrographs with 128,520 particles were used, for the GDH reconstruction 2,491 micrographs with 285,518 particles were used, and for apoferritin EMPIAR-10200 reconstruction 1,255 micrographs with 56,990 particles were used. The T20S and GDH micrographs have been uploaded to EMPIAR and EMDB as EMPIAR-10218 (1,173 micrographs) and EMDB-9233, and EMPIAR-10217 (2,491 micrographs) and EMDB-9203, respectively.

### File conversion

Locally acquired images and movies were recorded as 32-bit MRC files, the EMPIAR datasets were in TIFF format. IMOD (Kremer et al., 1996) was used to convert the images to various JPEG compression formats and EMAN2 (Tang et al., 2007) was used for all other image conversions.

## Results and Discussions

### Commonly used EM file formats for lossless storage

The electron microscopy (EM) field utilizes several file formats during data generation, ranging from vendor specific formats to standard processing formats e.g. MRC and TIFF (Cheng et al., 2015) As an example, the Leginon-Appion workflow utilizes a 32-bit MRC file format to save images from electron detectors. Concurrent with data collection on direct detector cameras, motion correction is run on-the-fly and two more images are created in addition to the unaligned sum—an aligned sum and a dose-weighted aligned sum—thereby providing researchers more flexibility in entering their processing workflow. However, these steps lead to a significant data expansion during ingestion because these steps result in an addition of tens of thousands of images generated per microscope per day.

One process that has been adopted to reduce the file size is to compress the raw movies after the motion correction. The overall potential compression ratio depends on whether the data has low redundancy to start with. We have used bzip2 compression for storage of our Gatan K2 dark corrected movies because of the reasonably high compression ratio, compatibility of the program across multiple platforms, and the ability to parallelize compression to keep up with data generation. Due to the nature of electron counting images from the K2, images may be compressed to ∼13-16% of the original file size. For example, a 50 frame K2 counting movie (3838 x 3710 pixels) in 32-bit MRC is 1,400 Mb, but reduces to 221 Mb when bzip2 compressed (Table 1). Since FalconIII counting images are natively provided as scaled 16-bit integer (and are gain and dark corrected) there is no advantage with upscaling to a 32-bit format. Also, they have a far less effective compression with a ratio roughly 75-80% of the original file size.

**Table 1:** Comparison of image file treatments and conversions.

| File format | K2 EC* (Mb) | K2 EC SR** (Mb) | K2 Summed image*** (Mb) |
| --- | --- | --- | --- |
| MRC | 1400 | 2700 | 55 |
| MRC with bz2<sup>a</sup> | 221 (15.8%) | 357 (13.2%) | 42 (76.4%) |
| Tiff | 1400 (100%) | 2700 (100%) | 55 (100%) |
| Tiff ZIP | 282 (20.1%) | 386 (14.3%) | 42 (76.4%) |
| Tiff LZW | 286 (20.4%) | 371 (13.7%) | 51 (92.7%) |
| PNG (8-bit) | 223 (15.9%) | 413 (15.3%) | 12 (21.8%) |
| JPEG000 (most loss) | 8.5 (0.6%) | 33 (1.2%) | 0.2 (0.4%) |
| JPEG050 | 188 (13.4%) | 33 (1.2%) | 2.7 (4.8%) |
| JPEG075 | 273 (19.5%) | 33 (1.2%) | 4.2 (7.6%) |
| JPEG100 (least loss) | 842 (60.1%) | 625 (23.1%) | 15 (27.3%) |

**Notes:** In parenthesis compression ratio from MRC to the target format for: EC electron counting (3838 x 3710 pixel image), SR super resolution (7676 x 7420 image), and summed image (3838 x 3710 pixel image). \*50 frames saved in 16-bit format, \*\*50 frames saved in 8-bit format, \*\*\*32-bit format.

**Notes:** In parenthesis compression ratio from MRC to the target format for: EC electron counting (3838 × 3710 pixel image), SR super resolution (7676 × 7420 image), and summed image (3838 × 3710 pixel image). *50 frames saved in 16-bit format, **50 frames saved in 8-bit format, ***32-bit format.

The TIFF file format is another possibility for high quality storage of image data. Similar to MRCs, TIFFs tend to be large in file size with 32-bit MRC and 32-bit TIFF image having identical file sizes. For TIFFs, the disk space may be reduced using native compression. The most common TIFF compressions are the Lempel-Ziv-Welch algorithm, or LZW, and ZIP compression. These strategies use codecs, or table-based lookup algorithms, that aim to reduce the size of the original image. Both LZW and ZIP are lossless compression methods and so will not degrade image quality.

For movie stacks, 32-bit compressed TIFFs have no data loss and compress to comparable file sizes as standard bzip2 compression. By using TIFF LZW the resultant file is 20% of the original file size in comparison to 16% of the file size using bzip2. (Table 1) The aligned sum image, however, only has a modest compression at 93% file size using TIFF LZW and 76% with bzip2. An advantage for TIFF files over bzip2 compression is that modern processing packages have TIFF LZW support, such that compressed images may be used directly for refinement. (Grant et al., 2018 and Zivanov et al., 2018). However, for the aligned sum images there is little to no file size savings using TIFF lossless compression, similar to the results of compression of FalconIII movie data.

### Data reduction through file format conversion

There are a number of additional image file formats often used in photography, but not so far for cryoEM image processing. Two commonly used photography file formats that support up to 24-bit color are PNG (Portable Network Graphics) and JPEG (Joint Photographic Experts Group). Electron micrographs typically are in grayscale so may be better suited to 8-bit file formats, which are also used in print media. PNG is a lossless file format and can utilize LZW compression similar to TIFF images. The disk savings when a 32-bit grayscale TIFF file is converted to an 8-bit grayscale PNG file is comparable with TIFF LZW with a compression of 16% for K2 counting movies, but there is also as a significant 22% compression for the aligned sum image, which is more than a 4-fold advantage in disk space savings over TIFF LZW (Table 1).

JPEG is a lossy file format that uses discrete cosine transform (DCT) to express a finite sequence of data points in terms of a sum of cosine functions (Hudson et al., 2017). The main reason that the JPEG format is avoided for archiving when images will be used for further processing is that they may lose quality each time they are compressed. JPEG has a range of compression ratios ranging from JPEG100 with the least amount of information loss (corresponding to 60% of the original file size for the frame stack and 27% for the aligned sum image) to JPEG000 with the most amount of information loss (corresponding to 0.4% of the original file size for the frame stack and 0.4% for the aligned sum image). For K2 super resolution images the compression ratio plateaued with a JPEG compression beyond JPEG075 most likely due to super resolution images having high redundancy with most of the data being zero (Table1).

To understand the effect these compression algorithms have on the images and subsequent processing, we compared EMPIAR-10200 apoferritin micrographs in 32-bit MRC, 8-bit PNG and JPEG075 (which was the compression level that super resolution images plateaued and also is the default compression ratio used in IMOD). For feedback on the progress of data collection, manual inspection of the micrograph and CTF estimation are two best practices utilized by several centers (Alewijnse et al., 2017). By visual inspection, qualitatively the micrographs in all three formats look similar if not identical (Figure 1A) even at 100X magnification (Figure 1B, top row). For a more quantitative measure CTFFind4 CTF estimation was performed on the three image formats and the package results with a confidence greater than 80% were obtained within 0.01Å. This result indicates that using PNG or JPEG images to monitor experiments, would provide sufficient feedback to have confidence in proceeding with data collection. Nevertheless, the act of converting a 32-bit image to an 8-bit image incurs precision loss. This can be seen by examining a sub-region of the image at the pixel level; the 32-bit to 8-bit conversion truncates the grey level to an integer which results in artifacts (Figure 1B, bottom row). These artifacts are known issues for JPEG file formats because the compression method is lossy meaning that some original image information is lost and cannot be restored, thereby potentially impacting image processing results.

**Figure 1.**
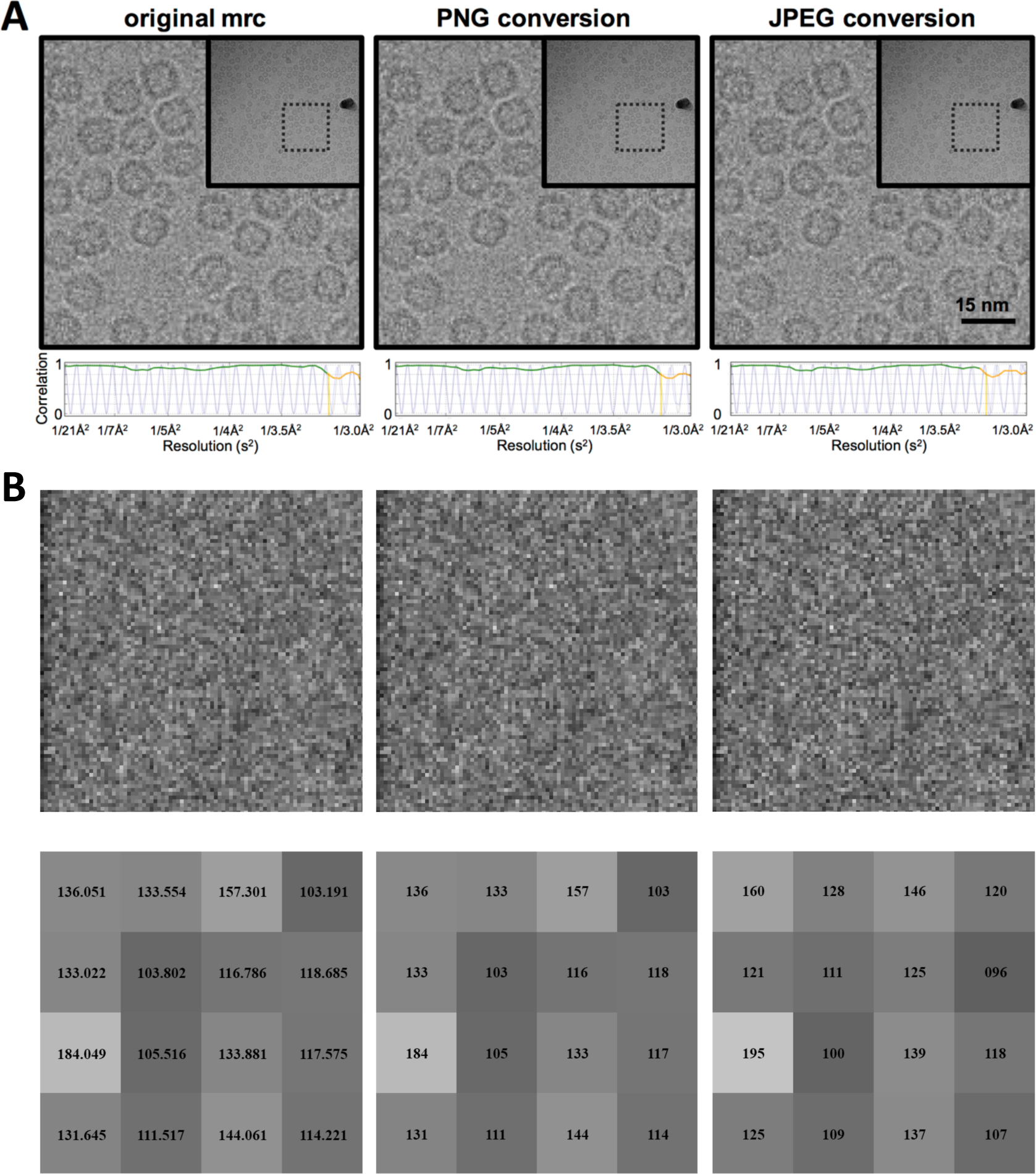
Effect of data reduction treatment on images. (A) Comparison of original MRC file, PNG lossless compressed image, and JPEG lossy compressed image at quality 75. (*Inset*) The original micrograph and region shown. Below each micrograph is a 1D plot of the CTF estimation by CTFFind4. In green is the 80% confidence level of the CTF estimation, which is at 3.18Å in the examples shown. (B) *Top*: Part of the image in 1A the original MRC format, after PNG compression and after JPEG quality 75 compression (100x magnification). *Bottom*: 4×4 selected pixel sub region area showing the pixel values before and after conversion.

### CryoEM reconstructions of JPEG images

Although there is a loss of data due to JPEG file conversion it is not obvious what the impact of the data loss is on the overall cryoEM workflow. To characterize this impact, we determined 3D reconstructions for standard test samples T20S and GDH using our standard workflow (Kim et al., 2018). These results showed that, 32-bit MRCs and 32-bit TIFF images may be used interchangeably for data processing. 3D refinement in RELION of a selected stack of 32-bit MRC particles following 2D classification provide maps at resolutions of FSC0.143 2.5 Å and 2.1 Å for T20S and GDH, respectively. When the particles were extracted from JPEG compressed micrographs and processed in the same way as the 32-bit MRC particles, we obtained similar reconstructions with identical FSC0.143 values (Figure 2a). Comparison of the unsharpened maps of GDH gave a cross correlation of 0.99. Examination of the maps shows that the overall structures are qualitatively quite similar, if not identical (Figure 2b).

**Figure 2.**
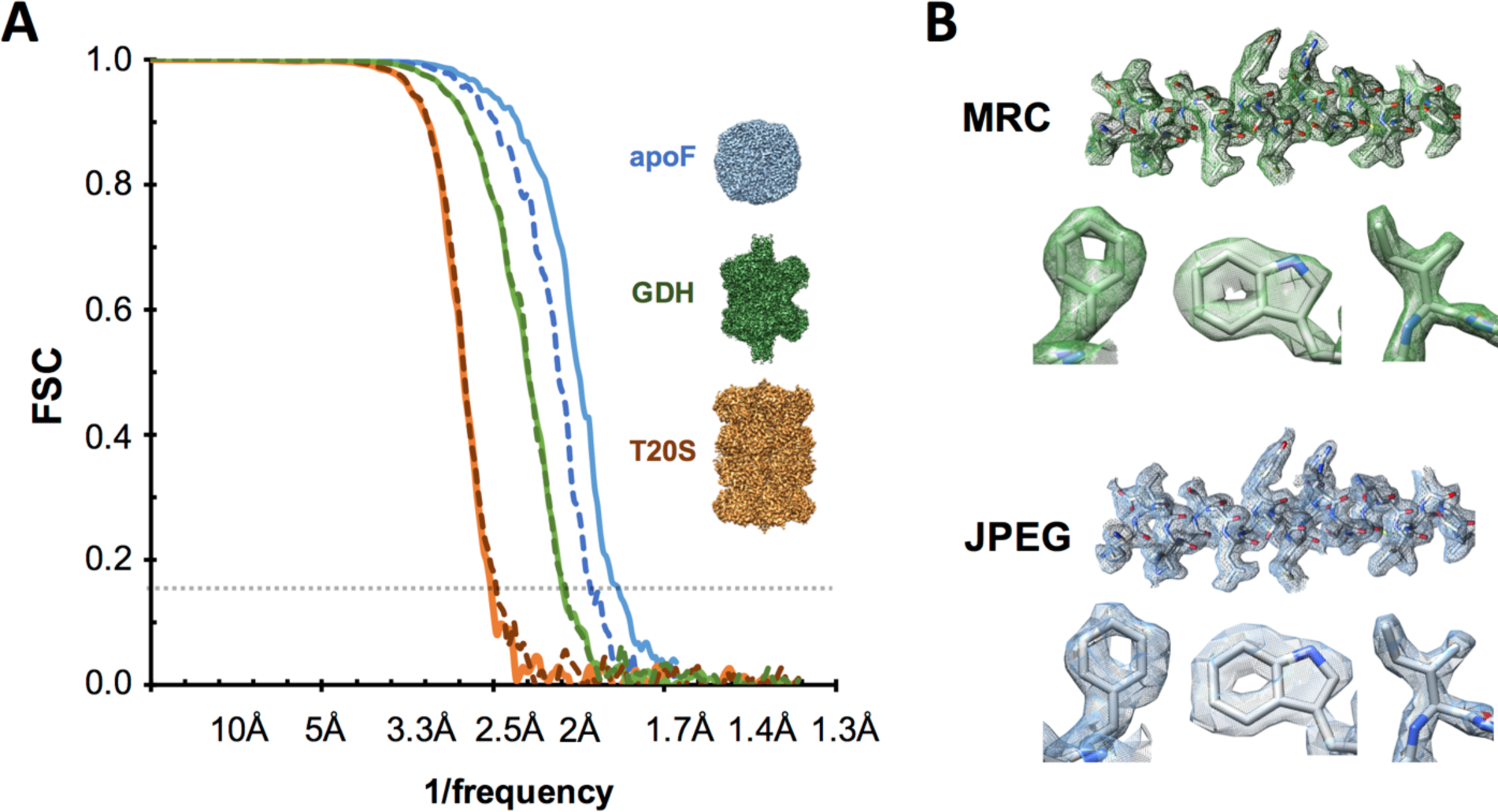
Comparison of untreated and treated data with test sets. (A) Fourier shell correlation of reconstructions with original MRC files (*solid line*) and lossy compressed JPEG quality 75 files (*dashed lines*) of T20S (*orange*) with 128,520 particles at FSC0.143 of 2.5Å, GDH (*green*) with 285,518 particles at FSC0.143 of 2.1Å, and EMPIAR-10200 apoferritin (*blue*) with 56,990 particles at FSC0.143 of 1.8Å. (B) 3D reconstruction comparison of selected areas of GDH reconstruction with original MRC (*green*) and JPEG compressed (*blue*) micrographs. Clockwise from the top is a representative α-helix in mesh, and representative parts of the maps: Ile β179, Trp β178 and Phe β399.

### Considerations when approaching Nyquist

Lossy compression on top of resampling may lead to unrecoverable loss of resolution. For the examples shown in Figure 2a the final resolutions at 2.5 Å and 2.1 Å are roughly half to two-thirds of the Nyquist frequency (pixel size 0.66 Å). We note that in Figure 2a the EMPIAR-10200 dataset is the only example where processing lossy images does not reach the same resolution as for lossless compressed images. The resolution obtained, FSC0.143 2.0 Å, is at ∼80% of the expect Nyquist value using JPEG compared to an FSC0.143 of 1.8 Å when using the MRC files (Figure 2a). In addition, CTF refinement and Bayesian polishing provided a further modest gain for a final FSC0.143 of 1.7 Å for MRC-extracted particles which is at 95% Nyquist.

To investigate this phenomenon further we used the cisTEM provided apoferritin test set (EMPIAR 10146 with a pixel size of 1.5Å and Nyquist frequency of 3.0Å) to determine how compression affects FSC0.143 resolution when close to Nyquist frequencies. Following the tutorial, reconstructions at Nyquist of 3.0 Å may be obtained with the raw uncompressed data. If the entire processing workflow (frame alignment, CTF estimation, particle picking, classification and refinement) is completed from PNG-converted movie stacks, then similar 3D reconstructions may be obtained as well with no apparent loss of resolution or quality of the resultant maps. Compression at JPEG100 (highest quality, least compression) produced a reconstruction within 0.1Å of the uncompressed data set with an FSC0.143 of 3.1Å. (Figure 3) We also tested the effect of lossy compression on the movie stack and a similar effect was noted. Increased compression with JPEG led to loss of resolution and impacted the ability to reproduce the original processing. By JPEG50 there is over a 10% decrease in resolution with FSC0.143 of 3.3 Å. Therefore, there will be a point where lossy compression will negatively impact the ability to recreate the experiment. This effect appears more severe the closer the experimental data is to the Nyquist frequency.

**Figure 3.**
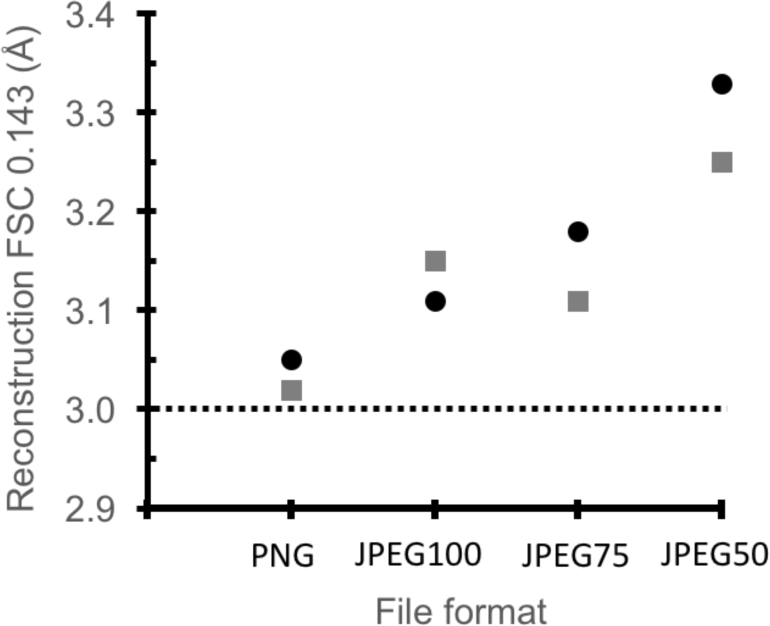
Apoferritin EMPIAR-10146 reconstruction FSC 0.143 versus file formats: PNG and different JPEG compression ratios. *Black circles* are aligned sum images and *grey squares* are movie stacks that were converted to PNG or various JPEG quality compressions before processing. The dotted horizontal line shows the uncompressed MRC reconstruction FSC0.143 of 3.0Å with a pixel size of 1.5Å.

## Conclusions

Ideally, the native format for all experimental data should be backed up and archived. At minimum, applying standard loss-less compression extends existing resources with a modest cost of computation time to restore the complete data set through decompression. If data generation greatly outpaces the ability to stage data for archival storage and impacts the ability to ingest new data, then data treatments may be utilized with no apparent impact on the final experiment outcome. We show that up to thirteen-fold data reduction by conversion to a lossy image file format (e.g. JPEG) for archival storage may be used to reproduce similar quality reconstructions as untreated data to at least 2 Å, pending the data is not Nyquist limited. JPEG utilizes DCT as one of the steps, discarding higher resolution frequencies as one of the lossy steps, with the level deciding how much is thrown away. Depending on compression level the treatment is only affecting data a certain distance from Nyquist. Lower resolution rings seem completely unaffected by the compression (as one would hope).

More modest approaches may be taken to reduce the potential data loss during these treatments, while maintaining the benefit of reduced storage requirements. Pending high data quality, modern refinement packages can process to Nyquist frequencies. Given there a r^2^ relationship for amount data in a resolution ring, discarding the outer rings provides substantial savings in space without losing quality, as long as there truly is no data in those rings. Therefore, cryoEM data may be binned or resampled at a pixel size half of the resolution based upon the final FSC_0.5_ or FSC0.143 value. Depending on the original image pixel size this may be a significant gain. These procedures do not obviate the need for backup and disaster recovery plans, but one can make informed decisions on how much data loss is acceptable for the gains in resource efficiencies.

## Acknowledgements

The work presented here was performed at the Simons Electron Microscopy Center and National Resource for Automated Molecular Microscopy located at the New York Structural Biology Center, supported by grants from the Simons Foundation (SF349247), NYSTAR, and the NIH National Institute of General Medical Sciences (GM103310) with additional support from Agouron Institute (F00316), NIH (OD019994) and NIH (RR029300).

